# Ecology of methyl-coenzyme M reductase encoding Thermoproteota

**DOI:** 10.1101/2025.10.16.682713

**Authors:** Zackary J. Jay, Matthew Kellom, Emiley Eloe-Fadrosh, Roland Hatzenpichler

**Affiliations:** Department of Chemistry and Biochemistry, Thermal Biology Institute, Montana State University, Bozeman, Montana, United States of America; Department of Energy Joint Genome Institute, Berkeley, California, United States of America; Department of Microbiology and Cell Biology, Center for Biofilm Engineering, Montana State University, Bozeman, Montana, United States of America

**Keywords:** archaea, carbon cycling, climate change, diversity, ecophysiology, MCR, methane, methanogens

## Abstract

The recent demonstration that members of at least three classes of archaea affiliated with the Thermoproteota superphylum are involved in the production of the climate-active gas methane has sparked discussions into how well we understand the diversity of methanogens. Here we show that members of all three of these lineages, as well as several other, yet uncultured and physiologically uncharacterized groups within the Thermoproteota that encode the key enzyme of anaerobic methane cycling, methyl-coenzyme M reductase (MCR), are widely distributed in anoxic ecosystems. We postulate that the taxonomic, metabolic, and ecological diversity of MCR-encoding Thermoproteota is poorly understood, and that the contribution of methylotrophic and thermoproteotal methanogenesis to methane production is largely unknown. We hypothesize that thermoproteotal methanogens contribute, potentially substantially, to methane emissions in many anoxic environments that harbor methylated precursors, including wetlands, sediments, peat, rice paddies, wastewater sludge, and geothermal systems. We highlight the necessity to experimentally test the (eco)physiology of these widely distributed archaea using both culture-dependent (*in vitro*) and culture-independent (*in situ*) approaches to assess their potential contribution to methane emissions. Last, we stress the importance of remaining agnostic about the physiology of MCR-encoding Thermoproteota in the absence of experimental data because most of these archaea also carry the genetic potential to grow non-methanogenically.

## Introduction

The first methanogens were cultured nearly a century ago^1^ and since then all cultured methanogens affiliated with only one of the four archaeal superphyla, the Euryarchaeota. However, over the last decade, methyl-coenzyme M reductase (MCR) and other methanogenesis marker genes were discovered through metagenomics on ∼250 environmentally derived metagenome-assembled genomes (MAGs) that do not affiliate with the Euryarchaeota. This led to the hypothesis that several lineages within the archaeal superphyla Thermoproteota and Asgardarchaeota might also engage in anaerobic methane cycling^2-19^. Until recently, all predictions on methanogenesis outside the Euryarchaeota lacked experimental validation. In 2024, three jointly published studies demonstrated that archaea affiliated with the thermoproteotal classes Methanosuratincolia^15,20^ and Korarchaiea^21^ were methyl-reducing methanogens. More recently, methyl-dismutating methanogenesis was demonstrated in a member of the thermoproteotal class Methanonezhaarchaeia^22^. Additionally, several other studies established that diverse archaea live by alkane oxidation^23-26^ and that a novel genus with the Euryarchaeota, *Ca*. Methanoglobus, contains methanogens^27-29^.

While these discoveries do not call into question the rate at which methane is released into the atmosphere, they demonstrate that we do not fully appreciate *which* organisms are involved in methane production. This is problematic because different methanogens likely vary in their metabolic profiles and ecological niches (*e*.*g*., which substrates they use, how they respond to environmental change) and in how they interact with other microorganisms. Thus, which specific methanogens are present and active in an ecosystem will have important consequences for ecosystem functioning – including methane emissions – competition for resources, impact on partner microbes, and resilience to change. A first step towards an understanding of the contribution of MCR-encoding Thermoproteota to methane production in the environment is to understand their ecology.

Here, we review the distribution of Thermoproteotal *mcrA* genes (encoding subunit alpha of MCR) in publicly available metagenomic and metatranscriptomic datasets on the Integrated Microbial Genomes and Metagenomes (IMG/M) and NCBI’s Short Read Archive (SRA) databases. Our goal is to highlight that the field has hitherto overlooked the potential contribution of these organisms to methane emissions and identify targets for future research.

### Group II Mcr sequences are near-exclusively found in the Thermoproteota

We focus our attention on “group II” (*sensu* Wang *et al*.^11^) *mcrA* sequences which, according to available data, are restricted to members of the Thermoproteota superphylum, with one exception. Members of the euryarchaeotal genus *Ca*. Methanoglobus likely have horizontally acquired their *mcr* genes and methanogenic potential^27^ from a member of the Thermoproteota with which they sometimes share their hot spring habitats^30^. By contrast, group I *mcrA* sequences are found exclusively in the Euryarchaeota. Group III sequences, which encode alkyl-coenzyme M reductases – distant relatives of MCR that use ethane, butane and longer-chain alkanes as substrates – have so far only been found in members of the Euryarchaeota as well as some MAGs affiliated with the Bathyarchaeia, Asgardarchaeota, and Methanosuratincolia^11,12^.

### Environmental distribution of *mcr* genes affiliated with the Thermoproteota

To better understand the diversity and distribution of MCR-encoding Thermoproteota in the environment, sequence searches were performed on unrestricted and publicly available datasets available from IMG/M (12.34 Tb as of April 2024) and SRA (1.75 Pb as of end 2021). From IMG, 589 metagenomes and 234 metatranscriptomes contained at least one group II *mcrA* sequence, compared to 322 metagenomes and 57 metatranscriptomes from NCBI-SRA. Samples had been collected from diverse habitats, including freshwater sediments, hot spring sediments, hot vents or seeps, oily sediments, coal beds, paddy soils, peat soils, marine sediments, and wastewater effluent or landfills (**Figs. 1-2, Table 1, SOI Datasets 1-2, Table S1**). Phylogenetic analysis, including reference McrA sequences (SOI Dataset 3), revealed distinctive clades corresponding to different taxa within the superphylum Thermoproteota, including Aukarchaeales, Bathyarchaeia, Methanoglobus, Methanonezhaarchaeia, Methanosuratincolia, Nitrososphaerota, and Korarchaeia (Supplementary Figure 1).

**Table 1.** Number of group II McrA/*mcrA* sequences detected in metagenomes (MG) or metatranscriptomes (MT) from either IMG or NCBI SRA databases, organized by taxa and generalized habitat. Numbers in brackets indicate the maximum PebbleScout score of the SRA count reported, with a minimum threshold score of 20 (scale = 20 - 100). The IMG search included 18,010 MGs and 5,302 MTs samples. The SRA search included 3,225,299 MGs and 62,103 MTs samples.

| Habitat | Taxa | JGI IMG |  | NCBI SRA |  |
| --- | --- | --- | --- | --- | --- |
|  |  | MG | MT | MG | MT |
| Freshwater sediment | Methanosuratincola | 138 | 83 | 33 [67.4] | 9 [100] |
|  | Methanomethylicus | 33 | 31 | 82 [100] | 23 [94.1] |
|  | Methanomethylovorus | 2 |  |  |  |
|  | Methanomethyloarchaeum |  | 1 |  |  |
|  | Bathyarchaeales | 48 | 11 | 3 [66.4] |  |
|  | Nitrososphaerota | 74 |  | 1 [89.5] |  |
|  | Korarchaeales | 1 |  |  |  |
| Peat soil | Methanosuratincola | 90 | 56 | 7 [57.0] |  |
| Hot spring | Methanosuratincola | 26 | 22 | 9 [41.0] |  |
|  | Methanomethylovorus | 41 | 2 | 9 [40.8] |  |
|  | Methanomethyloarchaeum | 27 |  | 1 [47.8] |  |
|  | Methanonezhaarchaeales | 6 |  |  |  |
|  | Nitrososphaerota |  |  | 3 [42.6] |  |
|  | Methanoglobus | 35 |  | 3 [24.0] |  |
|  | Korarchaeales | 12 |  | 3 [52.2] |  |
| Paddy soil | Methanosuratincola |  |  | 4 [48.7] | 2 [39.5] |
|  | Methanomethylicus |  |  | 2 [46.5] | 2 [43.1] |
| Wastewater, landfill | Methanosuratincola | 1 |  | 15 [47.5] |  |
|  | Methanomethylicus | 25 | 28 | 79 [100] | 15 [95.0] |
| Hot vent/seep | Aukarchaeales | 17 |  | 12 [82.6] |  |
|  | Methanonezhaarchaeales | 13 |  | 2 [45.9] |  |
|  | Methanoglobus |  |  | 1 [24.0] |  |
| Oil/gas | Methanosuratincola |  |  | 8 [35.6] | 2 [31.9] |
|  | Methanomethylicus |  |  | 9 [100] |  |
|  | Methanomethylovorus |  |  | 3 [22.5] | 3 [23.7] |
|  | Bathyarchaeales |  |  | 2 [28.2] |  |
|  | Methanoglobus |  |  | 1 [25.0] |  |
| Marine | Methanosuratincola |  |  | 4 [32.8] |  |
|  | Methanomethylicus |  |  | 5 [25.3] |  |
|  | Bathyarchaeales |  |  | 8 [32.8] |  |
|  | Methanonezhaarchaeales |  |  | 1 [31.4] |  |
|  | Korarchaeales |  |  | 4 [23.6] |  |
| Other | Methanosuratincola (Biocathode) |  |  | 1 [27.1] | 1 [24.1] |
|  | Methanomethylicus (Insect Gut) |  |  | 1 [21.8] |  |
|  | Methanomethylovorus (Book) |  |  | 4 [25.6] |  |
|  | Nitrososphaerota (Soil) |  |  | 1 [30.7] |  |

**Figure 1.**
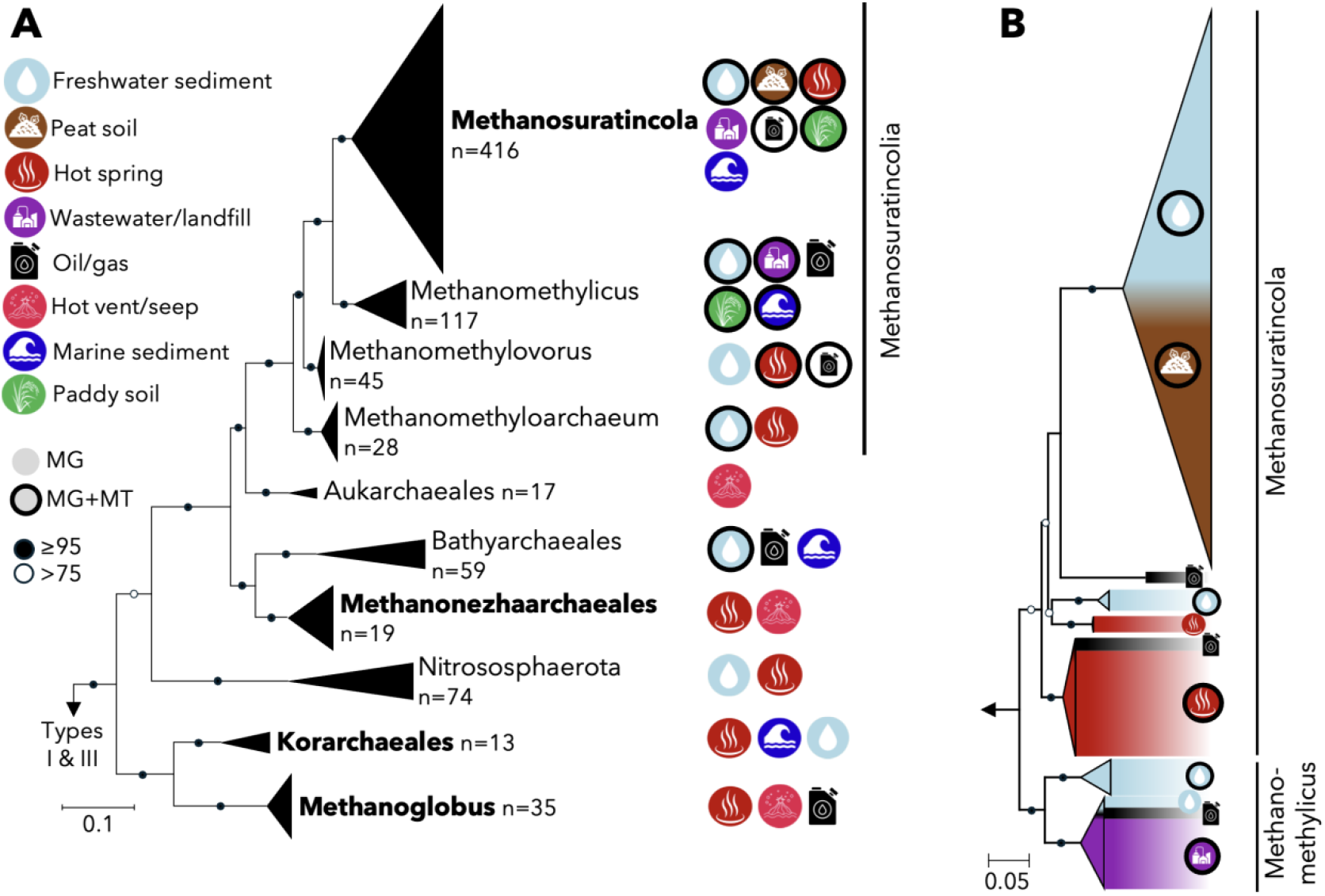
Phylogenetic analysis and habitat distribution of (**A**) all group II McrA sequences and (**B**) only Methanosuratincola- and Methanomethylicus-like group II McrA sequences. In both **A** and **B**, phylogenetic trees include reference and IMG sequences ≥200 amino acids. In **A**, sample counts only refer to the number of IMG protein sequences identified as SRA hits were not assembled. Habitat labels include additional habitats identified from the SRA. Circled icons specify that sequences were identified in both metagenomes (MG) and metatranscriptomes (MT). Taxa in bold indicate cultivated representatives. Habitat labels in **B** correlate only with reference and IMG sequences included in the tree. Refer to **Table 1** for sequence counts by database, dataset type, and habitat. Refer to **SOI Datasets 1 & 3** for IMG and reference group II McrA sequences, respectively. IMG sequences ≥200 amino acids, together with reference sequences (SOI Dataset 3), were phylogenetically analyzed with IQtree2^48^ (v. 2.0.6; LG+C60+F+G; 1000 ultrafast bootstrap) using trimAL^49^ (v1.4.rev22) trimmed (-gt 0.5) alignments generated with MAFFT-linsi^50^ (v7.522; -maxiterate 1000 -localpair) (Supplementary File 1).

**Figure 2.**
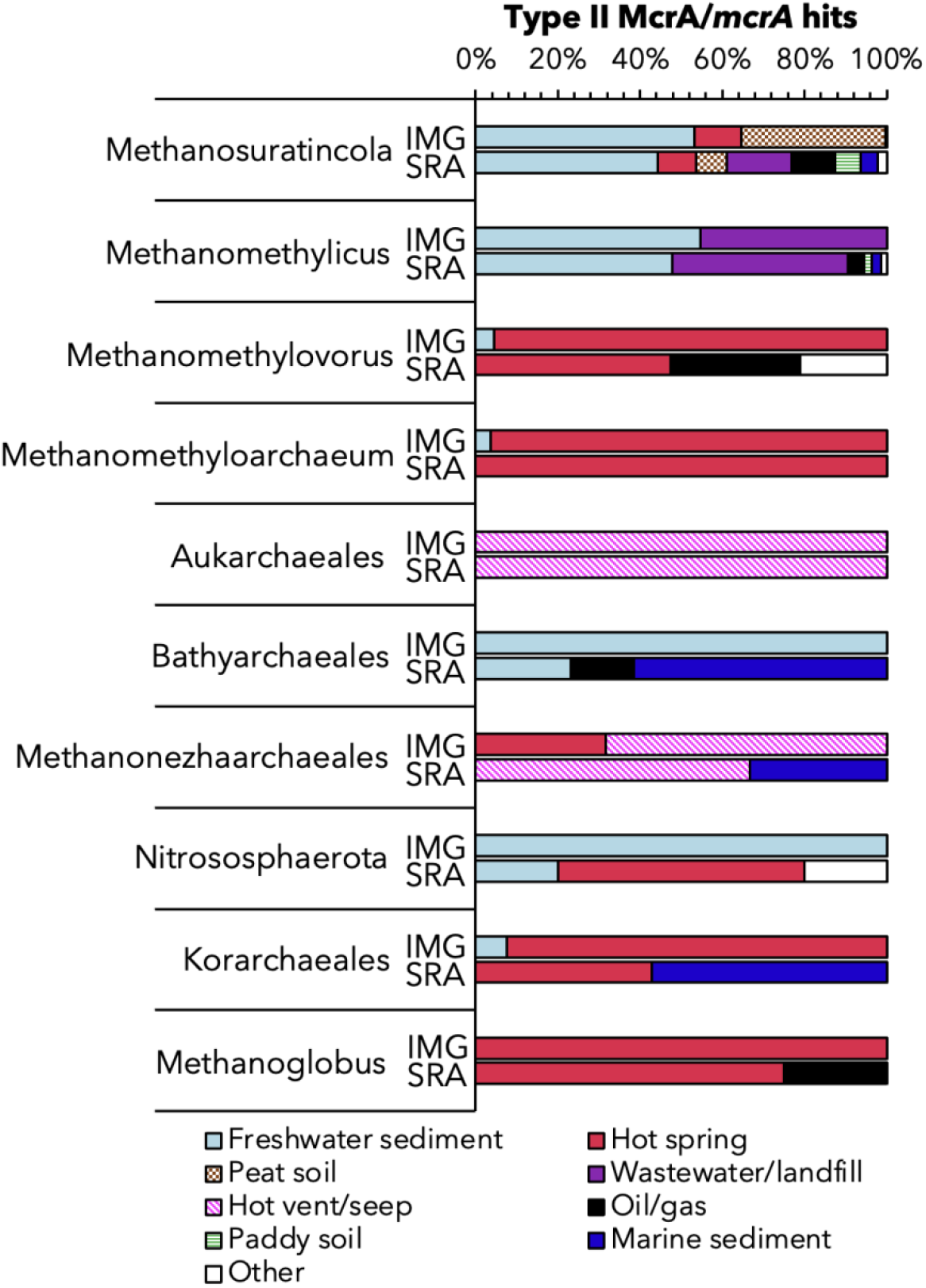
Relative proportion of habitats yielding group II McrA/*mcrA* hits from IMG or SRA, distributed by taxonomic assignment.

Interestingly, only a small proportion of IMG/M metagenome samples that contained group II *mcrA* also contained group I or group III sequences. Specifically, for the ‘Peat’ habitat category only 3 of the 77 group II containing metagenome samples also contained group I sequences. Other habitat types showed similar trends: 28 out of the 188 ‘Freshwater’ samples, 3 out of the 23 ‘Anaerobic digester’ samples, 5 out of the 58 ‘Hot Spring’ samples, and 3 out of the 10 ‘Hydrothermal Vent’ samples contained group I or III sequences; the remainder only contained group II *mcrA*’s.

Methanosuratincolia-like *mcrA* genes were the most abundant in both databases (IMG/M and NCBI-SRA), comprising 74 % of IMG and 88 % of SRA group II sequences. Of these, 51 % of IMG sequences were Methanosuratincola-like (versus 25 % in SRA) while 58 % SRA sequences were Methanomethylicus-like (versus 14 % in IMG). These differences may reflect disparities in search methodology or uneven dataset representation between databases. Indeed, both Methanosuratincola and Methanomethylicus genes were also the most widely distributed; they were found in both metagenomes and metatranscriptomes from many different habitats, including freshwater and hot spring sediments as well as paddy soils. Notably, all group II *mcrA* genes from peat metagenomes and metatranscriptomes were Methanosuratincola-like, whereas metagenomes and metatranscriptomes from wastewater or landfill were almost exclusively Methanomethylicus-like (**Fig. 1B**).

Other group II *mcrA* genes from Aukarchaeales, Bathyarchaeales, Methanoglobus, Methanonezhaarchaeaia, Nitrososphaerota, and Korarchaeales were much less abundant than Methanosuratincolia-like sequences but were nonetheless distributed in a variety of habitats (**Fig. 1A**). Methanonezhaarchaeia-, Korarchaeales- and Methanoglobus-like sequences were identified predominately in thermal habitats, although Korarchaeales-like sequences were also identified in mesophilic marine and freshwater sediments. In contrast, Bathyarchaeales- and Nitrosophaerota-like sequences were restricted to mesophilic sediments, although a few Nitrosophaerota-like sequences were also identified in hot spring metagenomes. The Aukarchaeales-like sequences were only identified in hot vent/seep metagenomes from Guaymas and Pescadero basins. This finding is likely a result of sampling bias. These two sites have received particular attention in recent years; we hypothesize that the comparatively large metagenomic datasets available for these sites^31-33^ allowed the detection of low-abundance archaea.

In summary, group II *mcrA* genes show taxonomic and habitat-specific associations, with Methanosuratincola-like sequences abundant in peat and freshwater sediments, Methanomethylicus-like sequences abundant in waterwater and landfills, and other taxa present at lower abundance in either thermal or mesophilic sediments.

### The environmental niche of *mcrA*-encoding Thermoproteota

With the exception of Methanonezhaarchaeia^11,22,34^, all currently discovered MCR-encoding Thermoproteota, share a key feature that distinguishes them from most euryarchaeotal methanogens: they lack both the tetrahydromethanopterin methyltransferase (Mtr) complex and the methyl-branch of the Wood-Ljungdahl (MBWL) pathway and thus depend on methylated substrates and substrate-specific methyltransferases^2,4-21^ to generate methane. The only reported cultivar within the Methanonezhaarchaeia^22^ represents an intriguing exception. Although this archaeon, like most MAGs affiliated with its class, encodes both Mtr and the MBWL pathway, it appears incapable of CO_2_-reducing methanogenesis and does not produce methane in the presence of H_2_. Under all conditions tested, it is an obligate methyl-dismutating methanogen^22^.

Before the discovery of MCR-encoding lineages within the Thermoproteota, methyl-dependent methanogenesis was thought to be relatively rare, with euryarchaeotal lineages exclusively using methylated precursors only having been occasionally observed^35-38^. However, methylated methanogenic substrates – including methanol and different methylamines – are widespread in nature and methyl-reducing methanogenesis is the dominant form of methanogenesis in anoxic marine, freshwater, and hypersaline sediments^37^.

A review of *in situ* measurements of methanol, mono-, di-, and tri-methyl amines in habitats similar to those where group II *mcrA* genes were identified, revealed a broad range of dissolved concentrations (**Table 2**). The highest concentrations of methanol were measured in freshwater sediments (2.0 mM) and peat soil samples (2.5 mM). Methylamine concentrations were not as high as methanol, although monomethylamine was measured as high as 9 µM in freshwater sediments (compared to 0.5 µM in marine samples), and concentrations of both dimethylamine and trimethylamine were measured as high as ∼50 µM in marine samples, compared to 1 and 2 µM in freshwater sediments. Methanol and methylamine concentrations have not been reported for thermal habitats (*e*.*g*., hot springs), where volatilization of these compounds likely makes measuring difficult, nor for oil-rich environments where they are presumably present within complex hydrocarbon mixtures.

**Table 2.** Reported dissolved concentrations of important methylated substrates measured in environments where group II *mcrA* sequences were identified. Please note that de Mesquita et al., (ref 37) contains many more relevant citations, which could not be listed here due to a limitation in citations for this article.

| Substrate | Environments | Concentrations | Citations |
| --- | --- | --- | --- |
| Methanol | Freshwater sediment | 40 nM – 1.92 mM | <sup>51</sup>, <sup>37</sup> |
|  | Peat soil | 480 µM – 2.6 mM | <sup>52</sup>, <sup>37</sup> |
|  | Marine sediment and ocean water | <12 nM – 150 µM | <sup>53</sup>, <sup>45</sup>, <sup>54</sup>, <sup>46</sup>, <sup>37</sup> |
|  | Paddy soil | ≤0.1 µM | <sup>37</sup> |
| Monomethylamine | Freshwater sediment | 150 nM – 8.9 µM | <sup>37</sup> |
|  | Marine sediment | 10 nM – 500 nM | <sup>37</sup>, <sup>55</sup> |
| Dimethylamine | Freshwater sediment | 400 nM – 900 nM | <sup>37</sup> |
|  | Marine sediment and ocean water | 0.8 nM – 48 µM | <sup>55</sup>, <sup>37</sup>, <sup>54</sup> |
| Trimethylamine | Freshwater sediment | 0.2 µM – 2.4 µM | <sup>37</sup> |
|  | Marine sediment and ocean water | 50 pM – 50 µM | <sup>37</sup>, <sup>55</sup>, <sup>54</sup> |

These data indicate that methylated substrates are widely available to group II McrA-encoding Thermoproteota, particularly Methanosuratincolia-like lineages, across diverse habitats. Why, then, have these organisms remained uncultivated for so long, even in habitats harboring traditional euryarchaeotal methanogens? One explanation could be that in environments dominated by traditional methanogens (*e*.*g*., freshwater sediments, peat, wastewater), MCR-encoding Thermoproteotaare less abundant and thus require deeper sequencing to be successfully assembled into a MAG. Consistent with this hypothesis, 91.6 % of IMG *Methanosuratincola*-like *mcrA* sequences had assembled coverages ≤20.2 reads per contig, and metagenomes and metatranscriptomes that generated up to 80 Gb of sequence accounted for 97.5 % of all the group II *mcrA* genes identified in IMG and SRA.

### Methanogens vs. MCR-encoding archaea

To date, four archaea within three classes of Thermoproteota (Methanosuratincolia, Methanonezhaarchaeia, and Korarchaeia) have been experimentally demonstrated to be capable of either methyl-reducing hydrogenotrophic^15,20,21^ or methyl-dismutating^22^ methanogenesis, and only one of them is available in pure culture^15^. There is, however, a larger number of MAGs representing other MCR-encoding lineages, including at least five other higher order taxa within three superphyla, *i*.*e*. Aukarchaeales^15^ Bathyarchaeia^2^ and Nitrososphaerota^34^ within the Thermoproteota, Helarchaeota^13^ within the Asgardarchaeota, and Hadesarchaea^4^ within the Euryarchaeota.

To the best of our knowledge, all group II *mcrA*-encoding MAGs as well as the genomes of the four available cultures encode energy conservation pathways other than methanogenesis. This is in stark contrast to group I methanogens within the Euryarchaeota, in which all but one representative – an engineered *Methanosarcina* sp. strain^39,40^ – appear to be obligate methanogens. For example, besides the potential to grow by reducing methyl-groups to methane, all MCR-encoding MAGs affiliated with the class Methanosuratincolia are genetically capable of conserving energy via amino acid or sugar fermentation^8,15,20,34,41^. Similarly, methanogenic members of the Korarchaeia might be able to grow by sulfite reduction to sulfide or via the anaerobic oxidation of methane^7,21^. Thus, MCR-encoding archaea within the Thermoproteota might not live exclusively by methanogenesis; rather, they may be better described as facultative methanogens. Depending on environmental conditions, substrate availability, and presence of other microbes that might compete for substrates, these archaea might adapt how they grow, which could give them a competitive advantage over obligate, euryarchaeotal methanogens. For example, methanogens must compete with other microbes for substrates, which can put them at thermodynamic disadvantage to other anaerobic energy-conservation pathways (*e*.*g*., sulfate reduction at high H_2_ levels). This has important implications for how we view MCR-encoding archaea within the Thermoproteota. However, so far, no experimental evidence for energy conservation pathways other than methanogenesis is available for any MCR-encoding Thermoproteota and tests for alternative substrates capable of sustaining growth were unsuccessful for the only isolate, *Methanosuratincola petrocarbonis*^15^.

Nevertheless, discovering *mcr* or other methanogenesis marker genes on a MAG does not directly implicate these organisms in methane cycling. In the absence of experimental data demonstrating their *in situ* or *in vitro* methanogenic activity – *e*.*g*., by demonstrating expression of *mcrA* mRNA^29^, single cell activity tests under methanogenic conditions^20,21^, or cultivation^15^ – we must remain agonistic to avoid making wrong conclusions and refer to them as MCR-encoding archaea rather than methanogens.

## Conclusion

Recovered thermoproteotal *mcrA* sequences were widely distributed and actively transcribed in diverse anoxic habitats. These included both natural and anthropogenic environments with high methane emissions^42^. The most diverse, widely distributed, and transcribed *mcrA* genes related to Methanosuratincolia methanogens were found in freshwater sediments – most importantly wetlands –, wastewater, and oil production sites. Together, these habitat types account for 65 % (365 million metric tons) of annual methane emissions^42^. Problematically, to the best of our knowledge, only two studies^5^ determined the activity of MCR-encoding Thermoproteota directly in their native habitat (by detection of *mcrA* mRNA via reverse transcription PCR^5^ and transcriptomics^43^). Furthermore, all available cultures of methanogenic Thermoproteota have been obtained from high temperature systems, *i*.*e*. three hot springs^20-22^ (64-77°C) and an oil reservoir^15^ (55°C). We are, however, unaware of their *in situ* activity and function of MCR-encoding Thermoproteota in all other ecosystems.

Based on these observations, we identify three key questions for future research:

1. Do MCR-encoding Thermoproteota grow by methanogenesis in their native ecosystems? To address this question researchers will need to elucidate their distribution, abundance, and methanogenic activity through targeted *in situ* and *in vitro*^29,30,44^ experiments as well as *in silico* analysis of existing sequencing datasets^56-58^. Such studies are needed to determine whether diverse MCR-encoding lineages indeed sustain themselves via methanogenesis.
2. How does methylotrophic methanogenesis contribute to methane production and emissions relative to CO_2_-reducing and acetoclastic pathways in these ecosystems, and what are the niche-differentiating factors driving the ecology and activity of thermoproteotal vs. euryarchaeotal methanogens? Answering these questions will require measuring the concentrations of methylated methanogenic precursors^37^, conducting substrate-specific rate measurements^45-47^, and collecting diverse environmental metadata.
3. Are thermoproteotal methanogens capable of using alternative energy conservation strategies (*e*.*g*., fermentation^8,15,20,34,41^, sulfite reduction^7,21^)? Using both culture-dependent and independent methods, researchers will need to test (meta)genomic hypotheses that MCR-encoding Thermoproteota might grow non-methanogenically. Confirmation of these hypotheses would fundamentally reshape our understanding of the biology of methanogens, because the mere detection of methanogenesis pathway genes would no longer implicate archaea in methane production, and it would highlight conditions under which non-methanogenic growth becomes favorable for these Thermoproteota.

We expect that the combined information gained from these experiments will provide a more holistic understanding of the biology and geochemistry of methanogenesis and, if methanogenic Thermoproteota can indeed shift between methanogenesis and non-methanogenic growth strategies, lead to novel strategies for mitigating methane emissions.

## Supporting information

McrA tree

## Author contributions

Conceptualization: R.H.

Formal analysis: Z.J.J., M.K., R.H.

Funding acquisition: R.H.

Investigation: Z.J.J., M.K., R.H.

Methodology: Z.J.J., M.K.

Project administration: R.H.

Supervision: R.H.

Visualization: Z.J.J., R.H.

Writing – original draft: Z.J.J., R.H.

Writing – review and editing: Z.J.J., M.K., E.E-F., R.H.

## Funding

This work was supported through award DE-SC0025661 by the U.S. DOE Biological and Environmental Research program to R.H. The work conducted by the U.S. Department of Energy Joint Genome Institute (https://ror.org/04xm1d337), a DOE Office of Science User Facility, is supported by the Office of Science of the U.S. Department of Energy operated under Contract No. DE-AC02-05CH11231. All opinions expressed in this paper are the authors’ and do not necessarily reflect the policies and views of the DOE.

## Declaration of competing interests

None.

## Supplementary information

### Supplementary files

**Methodology (below)**.

**Table S1 (excel file)**.

**Dataset 1 (excel file)**. IMG hits.

**Dataset 2 (excel file)**. SRA hits.

**Dataset 3 (excel file)**. Reference sequences.

**Supplementary Figure 1**. Un-collapsed version of the McrA tree shown in Figure 1.

### Methodology: querying public databases

Hidden Markov Models (HMMs), designed with reference Type II McrA sequences, were used to query the DOE-JGI IMG/M unrestricted and publicly available database (12.34 Tb as of Apr. 2024) at[ the National Energy Research Scientific Computing Center (NERSC). All recovered sequences (**SOI Dataset 1)** and references (**SOI Dataset 3**) were clustered with CD-HIT (v4.8.1; default settings) from 90–99 % amino acid identities in 1% increments. Manual curation of the clusters revealed clustering at 90 % amino acid identity was sufficient to separate sequences by taxa, although one cluster contained identifiable sequences from Methanosuratincola, Methanomethylovorus and Methanomethyloarchaeum at 90 % identity. These entries were separated by manual curation. IMG/M samples that contained group II sequences were manually inspected for the presence of group I and III sequences and confirmed through phylogenetic analysis (see Fig.1 caption).

To search NCBI SRA, query *mcrA* sequences were first identified by performing pairwise nucleotide comparisons (blastn) of sequences within each phylogenetic group and identifying divergent sequences, i.e., ≤90 % nucleotide identity. Nucleic acid sequences were submitted to NCBI’s PebbleScout (https://pebblescout.ncbi.nlm.nih.gov/) which queried all metagenomic and metatranscriptomic runs released in the public SRA before the end of 2021 (1.75 Pb). To minimize the inclusion of redundant IMG datasets, SRA results were excluded if the SRA Run (SRR) number matched those in IMG or if the provider was listed as ‘JGI’ or ‘Joint Genome Institute.’ This removed 60 % of the results, emphasizing the number of datasets provided by JGI to NCBI. Results were considered positive if the PebbleScout Score was ≥20, which included hits to unexpected habitats (i.e., ‘biocathode’, ‘insect gut’ and ‘book’ metagenomes) (**SOI Dataset 2**) but also may have included single hits more than once, especially considering the short fragment nature of the data. This, however, is probably rare as there is potential for more than one taxon to be found in any given habitat and the notable exclusiveness at this threshold of Methanosuratincola-like hits in peat and Methanomethylicus-like hits in wastewater/landfill (**Table 1**). It is important to note that datasets contributed after 2021 are not yet searchable by Pebblescout.

It is important to point out that most datasets on IMG/M and the SRA were created by research groups in the Global North and are biased towards sampling sites in the Northern hemisphere. This likely biased our understanding of microbial biogeography.

## Notes

### Competing Interest Statement

The authors have declared no competing interest.

### Summary of Updates

Updated manuscript text. No change to figures or tables.

