## Supplementary figures and images for "Ecology of methyl-coenzyme M reductase encoding Thermoproteota"

### McrA tree

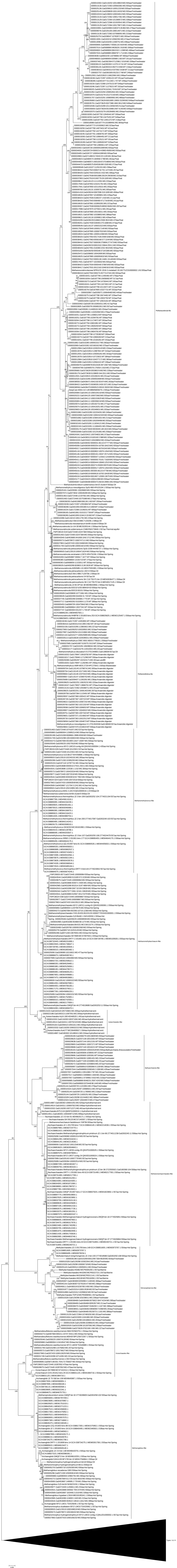
